# The Drosophila *Dopamine 2-like receptor D2R* (*Dop2R*) is required in the Blood Brain Barrier for male courtship

**DOI:** 10.1101/2022.11.22.516688

**Authors:** Cameron R. Love, Sumit Gautam, Chamala Lama, Nhu Hoa Le, Brigitte Dauwalder

## Abstract

The blood brain barrier (BBB) has the essential function to protect the brain from potentially hazardous molecules while also enabling controlled selective uptake. How these processes and signaling inside BBB cells control neuronal function is an intense area of interest. Signaling in the adult *Drosophila* BBB is required for normal male courtship behavior and relies on male-specific molecules in the BBB. Here we show that the dopamine receptor *D2R* is expressed in the BBB and is required in mature males for normal mating behavior. Conditional adult male knockdown of *D2R* in BBB cells causes courtship defects. The courtship defects observed in genetic *D2R* mutants can be rescued by expression of normal *D2R* specifically in the BBB of adult males. *Drosophila* BBB cells are glial cells. Our findings thus identify a specific glial function for the *DR2* receptor and dopamine signaling in the regulation of a complex behavior.

## Introduction

The neurotransmitter dopamine (DA) is evolutionarily highly conserved and has many described functions ^1^. In vertebrates, it has been shown to regulate motor function and locomotion, sexual behavior and response to drugs of abuse. In humans, it is associated with neurologic and psychiatric disorders such as Parkinsons and schizophrenia. As with other widely used signaling molecules, how limited and specific effects can be achieved is an important question. The availability of different receptors that vary in their distribution and employ different downstream signaling pathways is likely part of the answer. Several receptors have been identified that mediate DA signaling. Dopamine receptors are G-protein coupled seven-transmembrane receptors (GPCRs) that are classified into two broad classes: D1-like receptors (D1Rs) and D2-like receptors (D2Rs) based on their major described modes of signaling. D1Rs have been shown to signal through the G-protein G-alpha(s), leading to stimulation of adenylyl cyclase and to an increase in cAMP. Based on homology and functional analysis two D1-like receptors have been identified in *Drosophila,* Dop1R1 and Dop1R2 (DAMB) ^2–5^ with functions in classical conditioning and sleep.

D2R receptors are generally characterized by their ability to downregulate adenylyl cyclase and subsequent lowering of the second messenger cAMP. Human D2R has been associated with schizophrenia, addiction and Parkinson’s disease. D2 receptor agonists (e.g., bromocriptine) have proven efficacious in Parkinson’s disease, whereas antagonists at this receptor (e.g., haloperidol and chlorpromazine) are used commonly in the treatment of psychiatric illness ^6^. The *Drosophila* dopamine D2-like receptor D2R was isolated by homology with other dopamine receptors and is most similar to mammalian D2R. When expressed in a heterologous HEK392 cell system *Drosophila* D2R is activated by dopamine and other biogenic amines at micromolar concentrations, and is stimulated by the D2R agonist bromocriptine. Additional experiments in that system using PTX inhibition indicated that the receptor signals through Go/i as its mammalian counterparts do ^7^.

The *Drosophila* D2R gene codes for eight isoforms of the protein. Like its mammalian counterpart, a subset of them codes for a ‘short’ form of D2R with a shorter intracellular loop 3 than is found in the ‘long’ forms. This region is implicated in the receptor interaction with G-proteins. In mammals the long form has been found to be localized post-synaptically, while the short form was found presynaptically, functioning as an autoreceptor. Studies by Vickrey et al have shown that *Drosophila* D2R can function as an auto-receptor ^8^. *Drosophila* D2R has been shown to regulate locomotor activity and to play a role in learning and memory ^9–11^. Draper et al. found expression of D2R in neuronal and peripheral tissues such as portions of the gut and malpighian tubules. When they expressed a D2R RNA interference (D2R-RNAi) construct widely in the fly nervous system, they observed decreased locomotion. This defect was rescued when flies were fed bromocriptine, a D2R agonist ^9^.

The best-known D2R roles are in neuronal signaling. Little is known about potential glial roles of this receptor. In this study, we describe our finding that *Drosophila* D2R is expressed in the glial SPG cells of the blood brain barrier (BBB) where it plays an important role in the regulation of male courtship behavior. We have found that knockdown of D2R in the adult BBB leads to reduced male courtship. Courtship is similarly reduced in D2R mutants. Targeted BBB expression of the wildtype receptor in mature D2R-mutant males is capable of rescuing the defect, suggesting that D2R signaling in the BBB is required for the normal function of courtship circuits.

## Materials and Methods

### Fly stocks

*SPG-Gal4/TM3*^12^ was a gift from Roland Bainton (UCSF). *tubP-Gal80^ts^/CyO* and *tubP-Gal80t^s^/TM3,Sb* flies were a gift from Gregg Roman (University of Mississippi). D2R null mutant *Df(1)Dop2RΔ1, Dop2RΔ1* was a gift from Martin Schwärzel (Freie Universität Berlin). UAS-D2R RNAi line *y[1] sc[*] v[1] sev[21]; P{y[+t7.7] v[+t1.8]=TRiP.GL01057}attP2* (BL36824), *_w1118_ PBac{WH}Dop2R_f06521_* (BL 85250) D2R hypomorphic mutant; and *w*; P{PTT-GC}IndyYC0017/TM6C, Sb1 (Indy-GFP)* (BL 50860) were obtained from the Bloomington *Drosophila* stock center (https://bdsc.indiana.edu/).

D2R mutants were outcrossed for 10 generations with w^1118^(CS) flies (w^1118^ in a Cantonized background). The w^1118^ allele was exchanged for w^+^ in both outcrossed D2R hypomorph and null mutants by recombination.

### Behavioral assays

The courtship assay and activity assay were performed as previously described ^13^. Male courtship in *Drosophila melanogaster* consists of well-defined stereotyped behavioral steps that can easily be quantified in a courtship index (CI) ^14–16^. The CI is calculated as the fraction of time the male spends displaying any element of courtship behavior (orienting, following, wing extension, licking, attempted copulation, copulation) within a 10 minute observation period ^17^.

In short, males were placed in a plexiglass “mating wheel” (diameter 0.8 cm), together with a 2–4 hrs old *Canton-S* virgin female. Short-term activity assays were performed as previously described ^18^. Individual males were placed into the ‘‘mating wheel’’ containing a filter paper with a single line dividing the chamber in half. After 2–3 minutes of acclimation time, the number of times the male crossed the center line within the three-minute observation time was counted. Each graph represents sets of control and experimental genotypes that were grown, collected and aged in parallel. In each behavioral session, equal numbers of all genotypes were tested.

### Gal80^ts^ induction experiments

For conditional RNAi knockdown in mature males, *tubP-Gal80^ts^* carrying flies and control flies were raised at 18°C. Virgin males were collected at eclosion and kept in individual vials for 5–8 days at 18°C. Flies were then placed at 32°C for 48 hours for induction. Following induction, induced and uninduced flies were kept at 25°C overnight prior to courtship assays.

For RNAi knockdown during larval stages and reversal of knockdown in adulthood, food vials containing the fly crosses were kept at 18°C until wandering third instar was reached. Then half the crosses were shifted to 30°C until eclosion. Virgin males were collected at eclosion and kept in individual vials. They were kept at 30°C for one day and then transferred to 18°C for 6 days. They were then kept at 25°C for 1 day before being tested. Non-induced flies were kept at 18°C until 1 day before being tested when they were also transferred to 25°C.

### Bromocriptine experiments

Flies were reared at RT and transferred to Formula 4-24 Instant *Drosophila* Medium (Carolina), # 173200, containing DMSO or DMSO with 1 mM bromocriptine one day before the courtship assay.

### Generation of D2R transgene encoding V5-tagged D2R-RH

D2R-RH, a full length D2R mRNA isoform was synthesized by Invitrogen with the sequence coding for V5 (GKPIPNPLLGLDST) added at the carboxy terminus: GGT AAG CCT ATC CCT AAC CCT CTC CTC GGTCTC GAT TCT ACG STOP(TGA). The sequence was inserted into the transformation vector *pUAST* and transgenic flies generated by Rainbow Genetics, Inc.

### Immunohistochemistry

Immunohistochemistry on isolated brains was performed as described in Li et al. ^19^. The anti-D2R antibody was a gift from Isabelle Draper, Tufts Medical Center ^20^ and was used at 1:500 dilution. To visualize BBB cells, flies carrying *Indy-GFP* were used. *Indy-GFP* marks PG and SPG cells ^21^.

Antibodies used: Rabbit anti-D2R, 1:500 or 1:1000 (Draper ^20^); chicken anti-GFP (abcam ab13970), 1:500; Alexa Fluor 555 goat anti-rabbit (Invitrogen A21429) 1:200; Alexa Fluor 488 goat anti-chicken (Thermo Fisher Scientific A-11039).

Injection of 10kd Dextran-TR to assess the integrity of the BBB was performed as described in Hoxha et al. ^22^. Flies were subjected to the same experimental regimen as in behavioral experiments and control and experimental flies were assayed in parallel and imaged under identical conditions.

### Statistical Analysis

Two-way analysis of variance (ANOVA) was used to establish overall significance. Post hoc analysis for multiple comparisons was carried out with Tukey (HSD). P values < 0.05 were considered statistically significant. All statistical calculations were done using XLSTAT (Addinsoft, NY, NY) running on Microsoft Excel for Mac (version 16). All ±error bars are standard error of the mean (SEM).

### Data sharing

The data that support the findings of this study are available from the corresponding author upon reasonable request.

## Results

### D2R is present in the BBB

In a microarray analysis of isolated male and female BBB cells we identified D2R as a male-enriched BBB transcript ^23^. To confirm its presence in the BBB, we performed immunostaining of dissected brains using a D2R antibody created by Draper et al ^9^. *Drosophila* has an open circulatory system that distributes the hemolymph. It is separated from the brain by the BBB, two layers of glial cells that surround the brain like a tight cap. The outer layer of Perineurial Glia (PG) is thought to filter larger molecules. The inner Subperineurial Glia (SPG) are sealed with septate junctions and form the tight and selective blood brain barrier. Underneath the SPG lie the nuclei of the neuronal cortex ^24–26^. Figure 1 shows the presence of D2R staining around the brain, confirming the presence of D2R in the BBB. We compared D2R staining to staining seen with a *Indy-GFP* reporter line that marks both layers of the BBB (PG cells and SPG cells) ^27^. Double-staining of *Indy-GFP* brains with anti-GFP and anti-D2R shows that GFP and D2R colocalize on the basal side of the BBB, but not at the outer layer, indicating localization of D2R to the SPG cells (Figures 1a-c). For comparison, Figures 1d-f show double-staining in *Mdr-Gal4/ UAS-dsRed; Indy-GFP* flies. We have previously shown that *Mdr-Gal4* drives expression specifically in SPG cells ^23^. Like D2R, *Mdr-Gal4* driven dsRed is localized at the basal side. These data suggest that D2R is expressed specifically in the SPG cells of the BBB. This is consistent with the fact that the initial screen identifying D2R was conducted on isolated SPG cells ^23^.

**Figure 1.**
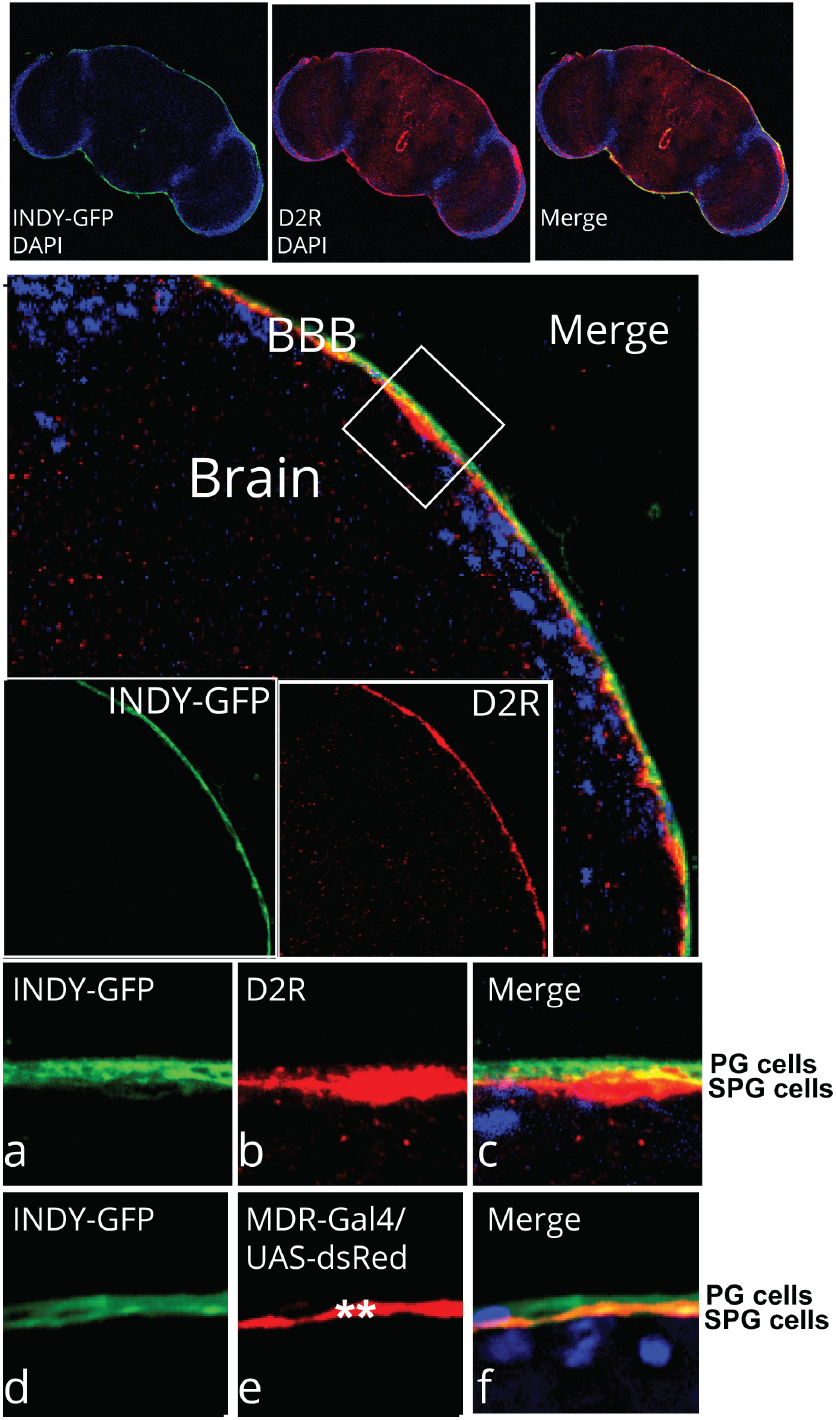
D2R is present in the SPG cells of the blood brain barrier. Isolated brains expressing Indy-GFP were stained with anti-D2R and anti-GFP antibodies. Indy-GFP is expressed in both layers of the blood brain barrier, the perineurial (SP) and the subperineurial (SPG) cells. Anti-D2R staining (red) can be seen in the BBB that surrounds the brain. It overlaps with *indy-GFP* staining (anti-GFP, green). Insets (63x magnification): Anti-GFP and D2R staining overlap on the basal side of the BBB (a-c). For comparison, double staining of SPG-specific *Mdr-Gal4* driven dsRed expression in SPG cells, marked by ** (anti-RFP, red) and indy-GFP (anti-GFP, green) is shown (d-f). Blue: DNA staining (DAPI).

### D2R knockdown in the BBB, but not neurons, reduces male courtship

To examine the possibility that *D2R* is required in the BBB for male courtship behavior, we conditionally expressed *D2R-RNAi* in the BBB of mature males using the *Gal4/Gal80^ts^/UAS* system ^28^ (Figure 2A). In this system, flies are reared and kept at 18°C. At this temperature, Gal4 is inhibited by Gal80^ts^ (“uninduced flies”). When animals are shifted to 32°C overnight, Gal80^ts^ is inactivated and Gal4 directs expression of *D2R-RNAi* (“induced flies”). Two different *SPG-cell-specific Gal4* drivers were used to direct expression, the previously described *SPG-Gal4*^12^ and our *Mdr65-Gal4* driver ^23^. The ATP binding cassette (ABC) transporter *Mdr65* has been shown to be specifically expressed in the SPG cells of the BBB, and *Mdr65-Gal4* reflects this expression ^23,29,30^. Control flies containing only one copy of the respective parental transgenes were grown, treated and tested in parallel to the knockdown flies. Male courtship in *Drosophila melanogaster* consists of well-defined stereotyped behavioral steps that can easily be quantified in a courtship index (CI) ^14–16^. The CI is calculated as the fraction of time the male spends displaying any element of courtship behavior (orienting, following, wing extension, licking, attempted copulation, copulation) within a 10 minute observation period ^17^. Flies that are continuously kept at 18°C, where Gal4 is inhibited by Gal80^ts^ and *D2R-RNAi* therefore not expressed, exhibit normal courtship, regardless of genotype. In contrast, when mature males were exposed to 32°C, males that expressed *D2R-RNAi* in the BBB had significantly reduced courtship. Reduction was observed with both the *SPG-Gal4* (Figure 2B; two-way ANOVA F(4,85)= 54.65, p<0.0001) or the *Mdr65-Gal4* driver (Figure 2C; two-way ANOVA F(3,65)= 8.77, p<0.0001). While courtship was reduced, the males were capable of performing all of the steps of courtship, but they did so with lower probability. To eliminate general sickness of the males as a cause for the reduced courtship, we performed a short-term activity assay ^18^. It assesses activity by counting the number of times a fly crosses a line drawn at the bottom of the courtship chamber. None of the flies with courtship defects showed reduced activity in comparison to the control flies (Figure 2D; two-way ANOVA F(3,96)= 0.99, p=0.400). These results indicate a physiological requirement for the D2R receptor in the BBB of adult males for normal courtship. To further explore D2R requirement we let *Mdr-Gal4/UAS-D2R-RNAi* and control flies develop at 18°C until they reached the wandering third instar stage. We then moved them to 30°C to induce *D2R-RNAi.* We collected individual males upon eclosion and continued 30°C incubation for half of the males until subjecting them to the courtship assay 4 days later. The other half were put to 18°C one day after eclosion for the next seven days (maturation at 18°C is slower than at 30°C) before being tested. At that temperature, Gal80^ts^ regains activity and *D2R-RNAi* expression ceases, allowing D2R function to resume. Experimental males that were continually kept at 30°C showed a courtship defect, while the courtship of control males was not affected by the higher temperature. Males that were moved back to 18°C after eclosion had normal courtship (Figure 2F; two-way ANOVA F(3,36)=76.14, p<0.0001). This result indicates that functional D2R in the BBB of adult males is sufficient for normal courtship, and that reduction of D2R during development does not cause an irreversible courtship defect. Together, these data support an adult physiological requirement for D2R in the BBB in the control of courtship.

**Figure 2.**
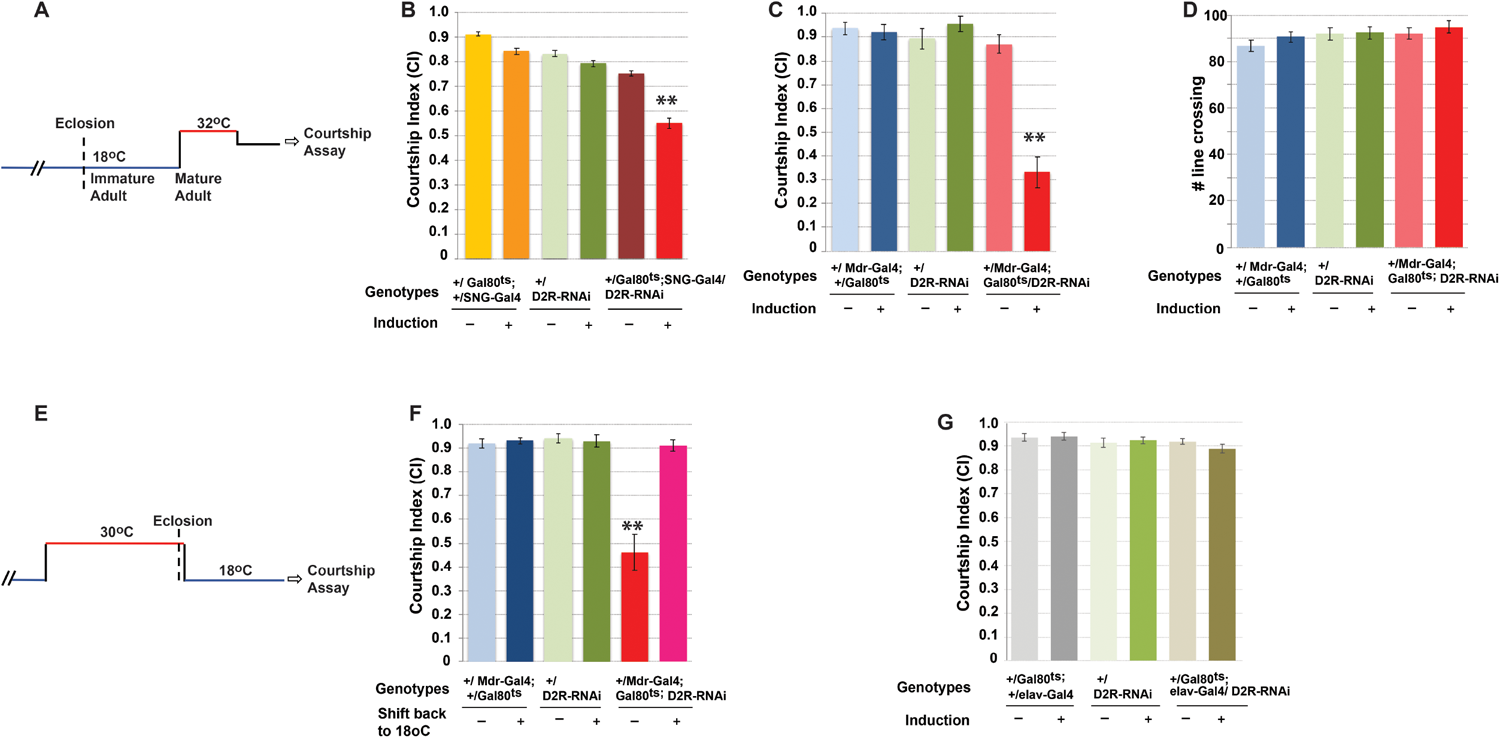
Conditional adult RNAi knockdown of D2R in SPG cells reduces male courtship. Conditional *D2R* RNA knockdown in the SPG cells of the BBB was performed. RNAi was expressed either in adult and mature males exclusively (A-D), or during development, with reversal during adult life (E, F). Graphs show the courtship index CI (fraction of time males spend courting during the observation period) ± SEM (B, C, F, G), or the performance of males in a control activity assay (# of line crossings ± SEM) (D) of the indicated genotypes. N= 20. Data were analyzed by ANOVA followed by Tukey multiple comparisons (p<0.05). Indices that are significantly different from the controls are marked by asterisks. *UAS-D2R-RNAi* expression is restricted by the presence of *tubP-Gal80^ts^* (abbreviated as *“Gal80^ts^”)* at 18°C (induction -). Placement of males at 32°C releases the Gal80 inhibition and leads to the expression of RNAi (induction +). Conditional expression of a *UAS-D2R-RNAi* transgene in mature males using two different SPG-specific drivers *(SNG-Gal4* (B) and *Mdr-Gal4* (C), respectively), significantly reduces male courtship in comparison to controls. The activity of the mutants as measured by number of line crossings is not different from control flies (D). Males with *D2R*-RNAi induction throughout development and adulthood had courtship defects, but when they were shifted back to reverse RNAi expression following eclosion, their courtship 5 days later was normal (F). Conditional knockdown of D2R in the neurons of mature males using the pan-neuronal driver *elav-Gal4* did not reduce their courtship (G).

Dopamine receptors are best known for their role in neuronal dopamine signaling. We next wanted to know whether knockdown of D2R in neurons would also affect courtship. We used a pan-neuronal driver *(elav-Gal4)* to conditionally express *D2R-RNAi* in adult neurons. As shown in Figure 2G, neuronal knockdown did not affect male courtship (two-way ANOVA F(5, 94)=0.66, p=0.652. These data suggest that signaling through D2R is important in the glial SPG cells of the BBB for courtship regulation but is not required in neurons.

### The courtship defects in D2R mutants can be rescued by expression of D2R in the BBB and by activation of the receptor

Next, we examined two D2R mutants (Figure 3). Dop2R^f06521^ is a hypomorph that results from the insertion of a Bac element in an intron. D2R^delta1^ is a null mutant that was generated by recombination of two Bac inserts and deletes the gene ^10^. Both mutants have reduced courtship (Figure 3A; two-way ANOVA F (3,96)=9.93, p<0.0001). None of the flies with courtship defects showed reduced activity in comparison to the control flies (Figure 3B; two-way ANOVA F(2,31)=0.08, p=0.923). These findings further confirm a role for D2R in male courtship behavior. To examine whether the mutant males might have compromised blood brain barriers, we performed a dye injection experiment that assesses the tightness of the barrier. Texas-Red (TR)-labelled 10 kD Dextran is injected into the abdomen from where it is distributed through the hemolymph in the open circulatory system. The molecule will accumulate at the BBB barrier and only diffuse into the brain when the barrier is compromised ^12,21^. As shown in Figure 3C, neither knockdown nor mutant flies showed defects in their barriers in comparison to the control.

**Figure 3.**
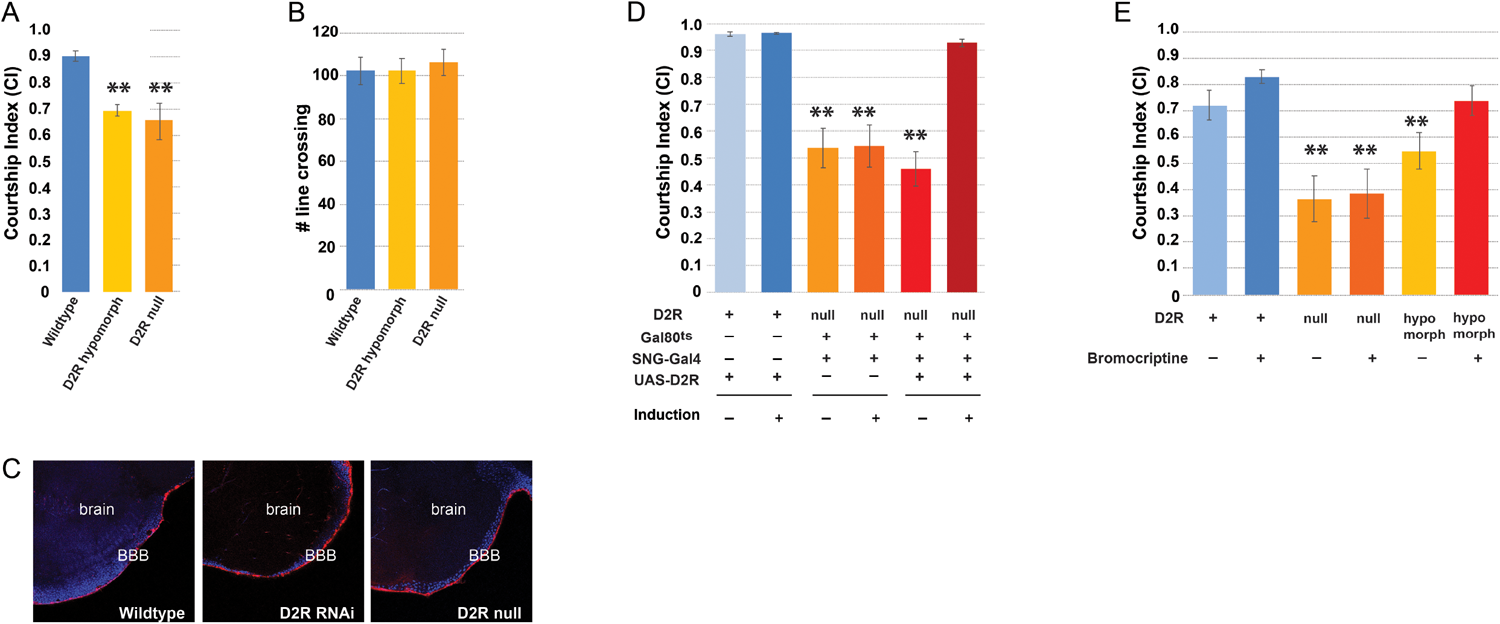
D2R mutants have courtship defects that can be rescued by expression of wildtype D2R in SPG cells or by a D2R agonist. *D2R* hypomorphic and null mutants have courtship defects (A), but normal activity (B). Conditional expression of wildtype *D2R* in SPG cells of adult D2R null mutants rescues courtship (D). Graphs show the courtship index CI (fraction of time males spend courting during the observation period) ± SEM (A, D, E), or the performance of males in a control activity assay (# of line crossings ± SEM) (B) of the indicated genotypes. N= 20. Data were analyzed by ANOVA followed by Tukey multiple comparisons (p<0.05). Indices that are significantly different from the controls are marked by asterisks. For rescue experiments, expression of full length *UAS-D2R* is restricted by the presence of *tubP-Gal80^ts^* (abbreviated as “*Gal80^ts^*”) at 18°C (induction -)(D). Placement of males at 32°C releases the Gal80 inhibition and leads to the expression of D2R (induction +). Feeding D2R mutants with Bromocriptine, a D2R agonist, rescues hypomorphic mutants, but not the null mutants (E). Blood–brain barrier integrity is not compromised in *D2R* mutant males (C). Experimental and controls flies were injected with 10 kDa TR-Dextran (red) and dye penetration into or exclusion from the brain was examined by confocal microscopy. The brain nuclei are stained with DAPI. A tight BBB is indicated by the demarcated red line on the surface of the brain indicating exclusion of TR-dextran from the brain. Flies were injected and processed in parallel and imaged with identical settings. Magnification = 63x.

We next tested whether expression of wildtype *D2R* in the BBB would rescue the courtship defect of the mutants. We created a *UAS-D2R* transgene and expressed it in the BBB of mature mutant males. As seen in Figure 3D, it was capable of rescuing the courtship defect of *D2R* mutant flies, in agreement with a requirement for *D2R* in the glial cells of the BBB for normal courtship (Two-way Anova: F (4,95) = 25.81, p< 0.0001). Bromocriptine is a well-described D2R agonist. Draper et al. have previously shown that *D2R* hypomorphs can be rescued by feeding the flies Bromocriptine. We fed adult D2R mutant males Bromocriptine and observed rescue of the courtship defect in the hypomorph, but not the null mutant (Figure 3E; two-way ANOVA F(4,75)=8.37, p<0.0001). *Dop2R^f06521^* hypomorphs still have low levels of D2R that can be activated by the agonist, whereas null mutants have no functional D2R receptor. This result suggests that rescue involves dopamine signaling through D2R. Together, our data suggest that active dopamine signaling in the BBB mediated by D2R is required for normal male courtship behavior. While most described roles of dopamine signaling are in neurons, we did not observe an effect when D2R was knocked down in neurons. Our data thus demonstrate a unique role for D2R in the glial SPG cells of the BBB in the control of a complex behavior.

## Discussion

The majority of what we know about dopamine involvement in courtship comes from studies that examined its roles in neurons. In *Drosophila,* dopamine and dopamine receptors other than *Dop2R* have previously been implicated in the regulation of courtship drive and the suppression of male-male courtship. Liu et al. have shown that males with (over-)activation of dopaminergic neurons show male-male courtship, without changes in male-female courtship ^31^. Similarly, a knockdown of the D1-like Dopamine receptor 1 *Dop1* (also called *DopR1 or DA1)* results in increased male-male courtship ^32^. dDA1 *(dumb)* mutant males were also shown to have reduced courtship drive as evidenced by a delay in courtship initiation and prolonged intervals between courtship bouts. This phenotype was rescued by expression of the wildtype receptor in mushroom bodies neurons ^33^. In contrast to *Dop1* mutants, knock-down of the D2-like receptor *D2R* or the *D1-like receptor 2 (DopR2, DAMB),* did not induce male–male courtship behavior ^32^. The role of dopamine in courtship drive and courtship motivation has been the focus of recent studies. Zhang et al have shown that the *D1-like receptor 2 (DopR2)* and dopaminergic circuitry underlie mating drive in males ^34^. The dopamine signal is transmitted through the *DopR2* receptor to P1 neurons, a group of *fruitless* expressing neurons, which integrate this signal with sensory information about the female. Similarly, dopaminergic activity in the superior medial protocerebrum (SMPa) reflects mating drive (influenced by previous matings and sperm depletion) and passes on that signal to P1 neurons through the *DopR2* receptor.

*D2R* had previously not been implicated in courtship regulation. Our data indicate that it is not required in neurons for courtship, as knockdown in neurons did not cause courtship defects. In contrast, we found that D2R has a specific role in the glial SPG cells of the adult BBB in the regulation of mating behavior. In mammals, some D2R isoforms have been found in glial cells in proximity to synapses. They play an important role as presynaptic autoreceptors, controlling dopamine levels by their involvement in dopamine turnover ^35^. Autoreceptor forms lack a portion of the second intracellular loop (‘D2R short forms”). Dopamine signaling has also been shown in astroglia of the prefrontal cortex ^36^ where application of a D2 agonist evoked elevated *Ca^2+^* levels. In contrast to the described glial autoreceptor forms of D2R, we used a ‘long’ full length D2R isoform in our rescue experiment, suggesting that postsynaptic dopamine signaling is responsible for D2R function in the glial SPG cells of the BBB.

In general, mammalian D2R has been shown to signal through Gi and inhibition of adenylyl cyclase, thus inhibiting cAMP signaling and PKA activation. When *Drosophila* D2R was expressed in mammalian HEK392 cells it was activated by dopamine (and to a lesser degree other biogenic amines), and signaling was inhibited by PTX. The S1 subunit of PTX from *B. pertussis* specifically ADP-ribosylates vertebrate G(i/o/t) proteins, resulting in their inability to bind to activated GPCRs ^37^. Flies do not have Gt, and their Gi lacks the site for ADP ribosylation. Thus, PTX in flies is specifically inhibiting Go ^38–40^. We have previously shown that expression of PTX in SPG cells or knockdown of Go in these cells reduces male courtship, identifying Go as a crucial mediator of BBB signaling for courtship. However, conditional inactivation of PKA signaling by dominant negative version of catalytic or regulatory subunits in the BBB had no effect on courtship ^22^. Furthermore, two well-known memory mutants with unregulated cAMP levels – *dunce* and *rutabaga*-have no courtship defects, suggesting that Go signaling in courtship is mediated by pathways other than cAMP signaling. Together these findings suggest that D2R signaling in the BBB is not mediated by cAMP. While mammalian D2R receptor signaling is thought to be mediated mainly by a decrease in cAMP-mediated signaling, alternative mediators have been described. Of particular interest is specific signaling through β-Arrestin. In addition to its involvement in termination of G protein mediated signaling, Arrestin is capable of initiating G-protein independent signaling cascades ^41^. One example is D2R signaling through interaction of Arrestin with Akt, PP2A and GSK ^42^. Interestingly, Peterson et al. have described biased signaling at the mouse dopamine D2 receptor, mediated alternatively by G-proteins or β-Arrestin ^43^. Their data suggest that D_2_R activates canonical MAP kinase activity through G proteins, whereas β-Arrestin may produce non-canonical ERK activity under conditions of enhanced β-Arrestin or kinase expression. Peterson at al. have further shown that different subsets of residues in the third intracellular loop of the receptor are required for the differential interactions. This is of particular interest since D2R is a prominent target for the treatment of schizophrenia, Alzheimer’s and depression, but many of these medications come with side effects. If these effects can be separated because they are based on the involvement of different downstream effector pathways, more specific therapeutics could be found. Whether biased D2R signaling in the BBB is involved the regulation of courtship is an interesting question for future studies.

The mammalian and *Drosophila* BBB are conserved in proteins that constitute the junctions that form the tight barrier. Similarly, functional proteins such as transporters or signaling molecules are highly conserved ^44^. Interestingly, profiling of the endothelial cells that form the mouse blood brain barrier has shown a “dopamine signaling signature”^45^ suggesting a conserved role for dopamine in these non-neuronal cells. How dopamine receptors and - signaling contributes to the function of this essential barrier remains to be elucidated. There is an increasing realization that the BBB has signaling roles beyond its barrier and transport function in the regulation of brain function. As we have shown here, signaling through the D2R receptor in the BBB is required for normal male courtship behavior, identifying a novel role for glial D2R and dopamine signaling in the control of a complex behavior.

### Conclusions

The Blood brain barrier (BBB) ensures that the brain is protected from harmful molecules, a function that is highly conserved across species. Besides this barrier role, there is increasing evidence that the BBB has functions in the regulation of brain processes. We have previously shown that in *Drosophila* it contains sex-specific factors that are required for normal male courtship behavior. Here we show that the dopamine receptor *D2R* is required in the BBB of mature males for normal mating behavior. These findings demonstrate that glial dopamine signaling is important for the regulation of a complex behavior.

